# Phenotype-driven transitions in regulatory network structure

**DOI:** 10.1101/142281

**Authors:** Megha Padi, John Quackenbush

## Abstract

Complex traits and diseases like human height or cancer are often not caused by a single mutation or genetic variant, but instead arise from multiple factors that together functionally perturb the underlying molecular network. Biological networks are known to be highly modular and contain dense “communities” of genes that carry out cellular processes, but these structures change between tissues, during development, and in disease. While many methods exist for inferring networks, we lack robust methods for quantifying changes in network structure. Here, we describe ALPACA (**A**Ltered **P**artitions **A**cross **C**ommunity **A**rchitectures), a method for comparing two genome-scale networks derived from different phenotypic states to identify condition-specific modules. In simulations, ALPACA leads to more nuanced, sensitive, and robust module discovery than currently available network comparison methods. We used ALPACA to compare transcriptional networks in three contexts: angiogenic and non-angiogenic subtypes of ovarian cancer, human fibroblasts expressing transforming viral oncogenes, and sexual dimorphism in human breast tissue. In each case, ALPACA identified modules enriched for processes relevant to the phenotype. For example, modules specific to angiogenic ovarian tumors were enriched for genes associated with blood vessel development, interferon signaling, and flavonoid biosynthesis. In comparing the modular structure of networks in female and male breast tissue, we found that female breast has distinct modules enriched for genes involved in estrogen receptor and ERK signaling. The functional relevance of these new modules indicate that not only does phenotypic change correlate with network structural changes, but also that ALPACA can identify such modules in complex networks.

**Significance statement:** Distinct phenotypes are often thought of in terms of unique patterns of gene expression. But the expression levels of genes and proteins are driven by networks of interacting elements, and changes in expression are driven by changes in the structure of the associated networks. Because of the size and complexity of these networks, identifying functionally significant changes in network topology has been an ongoing challenge. We describe a new method for comparing networks derived from related conditions, such as healthy and disease tissue, and identifying emergent modules associated with the phenotypic differences between the conditions. We show that this method can find both known and previously unreported pathways involved in three contexts: ovarian cancer, tumor viruses, and breast tissue development.

## INTRODUCTION

We tend to think of phenotypes as being characterized by differentially expressed genes or mutations in particular genes. However, the individual genes that show the greatest changes in expression in a phenotype do not tend to be drivers of that phenotype (1, 2). Despite the increasing power and depth of sequencing studies, identifying the causal mutations and Single Nucleotide Polymorphisms (SNPs) that are responsible for determining heritable traits and disease susceptibility remains challenging. Indeed, many studies have found thousands of genetic variants of small effect size contribute to common traits (3–5). It has become apparent that phenotypes are driven by complex regulatory interactions between multiple genes and variants that together define the state of the cell. Modeling these phenotypes requires that we have a clearer picture of how genes and proteins work together to perform normal cellular functions, and how remodeling the interactions between genes can cause changes in phenotype including disease.

In this context, we need to subtly shift our understanding and think of a phenotype as being defined by a network of interacting genes and gene products. Exploring the topology of such networks can provide important biological insight into phenotypic properties. For example, high-degree “hubs” in protein-protein interaction (PPI) networks are enriched for genes essential to growth (6). Biological networks are known to have modular structure and contain closely interacting groups of nodes, or “communities,” that work together to carry out cellular functions (7–9). There are many analytical and experimental methods for inferring network models associated with different phenotypic states (10–12). However, the most significant questions we can ask of biological networks – how networks differ from each other, and how differences in network structure relate to functional changes – remain largely unanswered.

Most analyses of so-called “differential networks” have focused on determining which edges are altered relative to a reference network (13). While the advantage of this approach is its simplicity, there are several issues that arise in such an edge-based analysis. First, biological network inference has a relatively high rate of false negatives due to noise in both the experimental data that are used and in the network inference methods themselves. Consequently, it can be difficult to determine whether the appearance or disappearance of a single edge is “real.” The uncertainty in the estimate of the difference between two edge weights is the sum of the uncertainties in each individual edge, which inflates noise in the final differential network. Second, the perturbed network will in general contain both positive and negative changes in edge weight relative to the reference network, and it is challenging to analyze and interpret a differential network with mixed signs. If we only consider the new edges associated with a phenotype, we would miss the functional effects of decreases in edge activity. Third, by focusing only on the altered edges and discarding common edges, the differential interactions are taken out of their functional context, making it challenging to connect them to global cellular changes. For example, adding or deleting ten scattered edges in a network may have very different consequences on the phenotype than would the same number of changes concentrated in a local functional neighborhood of the network.

One way to address these issues and find more robust differences between networks is to identify changes in groups of nodes, rather than in individual edges. Computational methods that have been developed to do this fall into several categories. First there are methods that evaluate differences in pre-specified network features, like user-defined gene sets, small regulatory motifs or global topological characteristics. For example, Gamberdella et al. evaluated the statistical significance of differences in co-expression of a user-defined gene set between two conditions (14). Similarly, the coXpress method defines clusters using co-expression in the reference condition, and tests for significant changes in each cluster under a new condition (15). Landeghem et al. developed a method for inferring the best differential network that contrasts two datasets, and Gill et al. and Danon et al. introduced new measures to test whether global modular structure and degree characteristics are different between two networks (16–18). However, these methods are limited to examining pre-defined gene modules and network features, and fail to take full advantage of the network structure. As such, they lack the ability to discover new pathways and network modules that functionally distinguish different phenotypes.

Other methods have been developed to discover *de novo* gene modules that differ between conditions. The DiffCoEx algorithm iteratively groups genes that are differentially co-expressed to find new modules (19, 20). Valcarcel et al. compared metabolite correlation networks to discover groups of metabolites that changed their correlation pattern between normal weight and obese mice (21). These methods are based on first computing the most differential edges and then grouping them together, which increases the uncertainty of each edge estimate and does not incorporate functional edges that are present in both conditions (13, 22), thus losing network context.

Another class of methods attempts to identify “active modules,” which are groups of genes that are differentially expressed in a particular disease or condition and also highly connected in a reference network, such as the protein-protein interaction network (23). However, the “active modules” framework only uses differential gene expression and so focuses on the nodes rather than accounting for changes in the strength of regulatory edges.

We present a new graph-based approach called <u>AL</u>tered <u>P</u>artitions <u>A</u>cross <u>C</u>ommunity <u>A</u>rchitectures (ALPACA) that compares two networks and identifies *de novo* the gene modules that arise in the networks as the phenotype changes. ALPACA is based on modularity maximization, a technique commonly used to find communities in a single graph. As applied previously, modularity is a measure of the observed edge density of the communities as compared to their expected density in a degree-matched random graph. Although this technique is powerful, it has a “resolution limit” because communities can only be identified if they are larger than the typical cluster size in random graph configurations (24). This lack of resolution is especially disadvantageous when studying transcriptional networks, which tend to have a dense and hierarchical structure, and whose functional units only become evident under different environmental conditions (25). A framework based on modularity maximization has been created to find common community structure among multiple networks (26), but the only way to detect differences is to apply modularity maximization to each network separately, followed by brute-force comparison of the two resulting community structures.

In ALPACA, we adapt the modularity framework to compare condition-specific networks to each other rather than to a random graph null model. We define a score called the “differential modularity” that compares the density of modules in the “perturbed” network to the expected density in a matched “baseline” network, allowing us to contrast, for example, networks from disease and healthy tissue samples and partition the nodes into optimal differential modules, without relying on predefined gene sets or pathways. In contrast to methods that simply cluster the most differential edges, ALPACA compares the full network structures active in each condition and reduces the noise from individual edges by estimating an aggregated null model. And because the null model is based on the properties of a known reference network rather than on a random graph, the usual “resolution limit” does not apply, and ALPACA can detect small disease modules otherwise hidden within larger regulatory programs associated with normal cellular functions.

To demonstrate the utility of ALPACA, we apply it to compare simulated networks, as well as transcriptional network pairs from non-angiogenic and angiogenic subtypes of ovarian cancer, normal human fibroblasts and fibroblasts expressing tumor virus oncogenes, and male and female breast tissue from the Genotype-Tissue Expression (GTEx) project. We find that ALPACA produces higher resolution and robustness than other network approaches and identifies modules enriched in biological processes relevant to the phenotypes we are comparing. Although we have focused on transcriptional networks, the framework we present here is mathematically general and could be applied to find the differences in modular structure between any two networks.

## RESULTS

### Modularity maximization and comparing community structures

Many methods for determining the community structure of a network are based on maximizing the modularity (27):

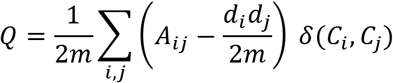

Here, *A_ij_* indicates the adjacency matrix of the network, *m* is the number of edges, *d_i_* is the degree of node *i*, and *C_i_* is the community assignment of node *i*. The modularity represents to what extent the proposed communities have more edges within them than expected in a randomly connected graph with the same degree properties; this null expectation is represented in the second term of the equation above. The modularity is optimized over the space of all possible partitions {*C*} and the value of *C_i_* corresponding to the maximum modularity then determines the community structure of the network. An exhaustive search is not possible for large networks, but many methods have been developed to find locally optimal community structure, including ones based on edge betweenness, label propagation, and random walks (27–29). The Louvain algorithm is a particularly efficient way to find high-quality local optima of the modularity function (30).

### Community comparison and edge subtraction

Having arrived at a pair of inferred networks corresponding to different phenotypic states, there are two straightforward ways to compare the community structures based on the modularity metric (Figure 1). One method, which we will call “community comparison,” consists of using modularity maximization to find the community structure for each network individually, and then finding the nodes that alter their community membership between the two networks. Another method, which we will call “edge subtraction,” is to compute the differences in the edge weights between the two networks, and then apply modularity maximization to the resulting subtracted weights.

**Figure 1.**
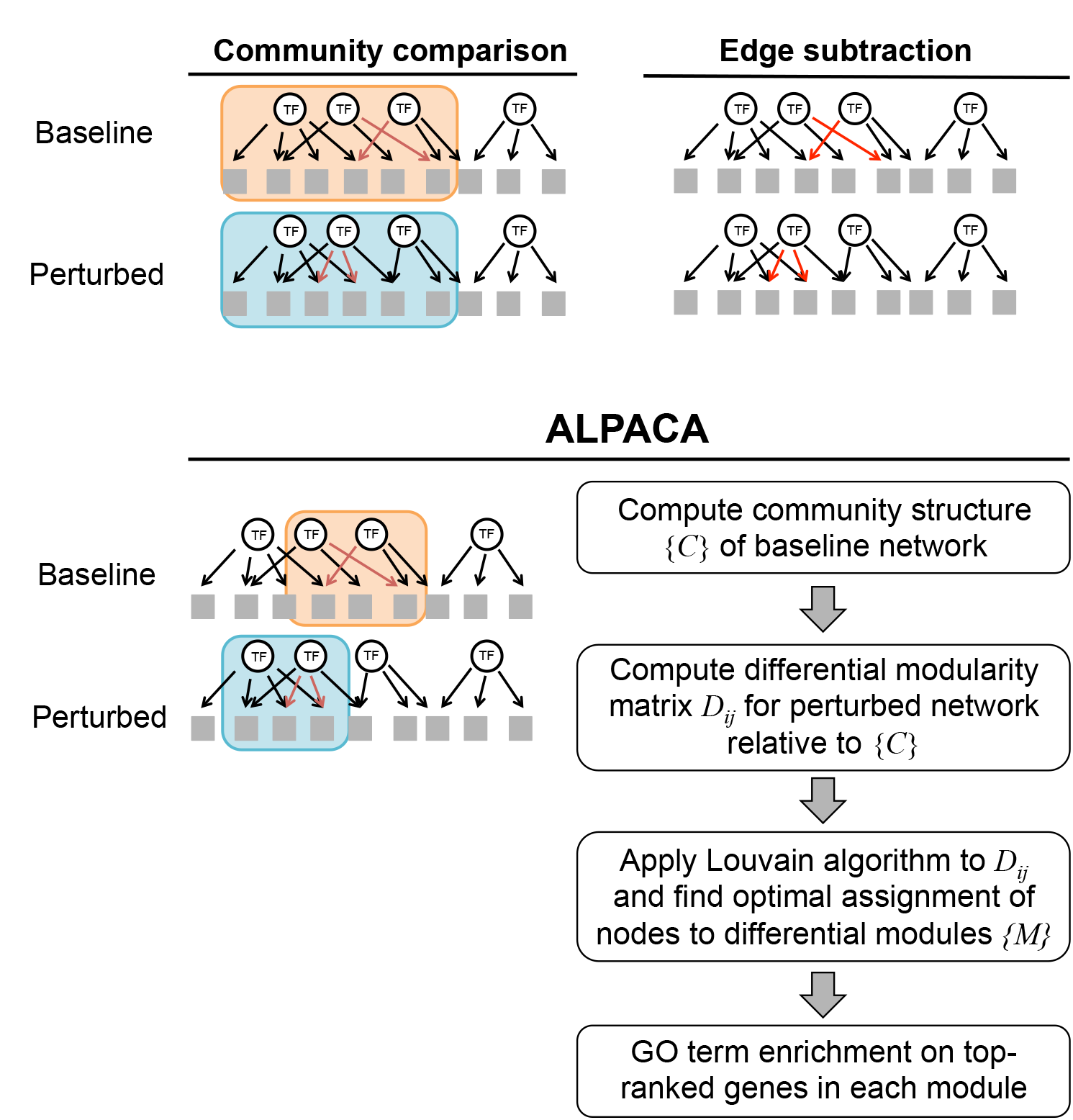
Methods to compare networks and find changes in modular structure. “Community comparison” identifies communities separately in each network and looks for nodes that change their community membership. “Edge subtraction” finds communities by subtracting the networks and finding communities in the resulting differential edges (red arrows). ALPACA looks for groups of genes that are more interconnected in the perturbed network than expected given the community structure of the baseline network. Flowchart shows the major steps in the implementation of ALPACA.

Both methods can detect large, dramatic changes in network structure. However, there are important differences in these methods. “Community comparison” is limited in its ability to detect structural changes smaller than the average community size in each individual network. In contrast, “edge subtraction” acts on the difference of the edge weights, which reduces the density of the network and increases the resolution, but this method is also more strongly affected by noise in the individual edges. Further, only positive edge weight differences can be used to run modularity maximization in the subtracted network, so edges that are lost are not appropriately accounted for; incorporating both positive and negative edge weight differences requires more complex techniques (31, 32).

### ALPACA: A new method for detecting changes in community structure

To overcome some of the limitations of the community comparison and edge subtraction methods, we developed ALPACA, a new algorithm based on modularity maximization. The unique aspect of ALPACA is that, rather than comparing edge distributions to a random null model, we compare edges of the “perturbed” network to a null model based on the “baseline” network to find differential gene modules between the two networks (Figure 1). ALPACA optimizes a new quantity called “differential modularity,” which we define as

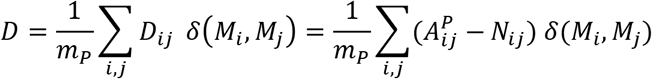

This score compares the number of edges in a module *M* in the perturbed network – whose adjacency matrix is given by 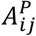 and total edge weight is *m_P_* – to the expected number of edges based on the pre-computed community structure {*C*} of the baseline network. Here, *N_ij_* is defined as

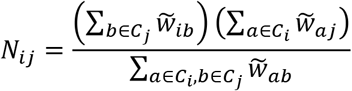

where *C_i_* is the community assignment of node *i* in the baseline network, and 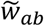 is the normalized weight of the edge between node *a* and node *b* in the baseline network: 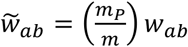. For the normalization, we have chosen to globally scale the edge weights of the baseline network so that the total matches *m^P^*, the sum of the edge weights in the perturbed network. This allows a fair comparison between two networks that could be derived from two datasets of differing quality or sample size and may have different global sensitivity properties. To identify the modules {*M*} that maximize the differential modularity, we use the following two-step procedure. First, we determine the community structure of the baseline network using established methods (9, 30). Second, we compute the differential modularity matrix *D_ij_* and apply the Louvain optimization algorithm to iteratively aggregate the nodes into modules (30).

Note that the equation above is presented in a form that applies to weighted bipartite networks, as we will be applying it to analyze transcription factor (TF)-gene interactions. It can be easily adapted to analyze other types of networks. More details about the implementation of all three methods – community comparison, edge subtraction, and differential modularity – are presented in the Materials and Methods section.

### Evaluating the performance of ALPACA on simulated networks

We reasoned that ALPACA would be more sensitive to small changes in modular structure than methods based on standard community detection, because the null model is computed using detailed properties of the baseline network rather than relying on random graphs. We also believed that ALPACA would be less sensitive to noise in individual edge weights than edge subtraction, because the null model is estimated by averaging over communities in the baseline network. We set out to test these properties in a setting that resembles real biological networks as much as possible, but where we have control over the changes in modularity.

To do this, we constructed a baseline network and then created new modules through the “addition” of new edges, resulting in a perturbed network. For the noiseless version of this simulation, we inferred a regulatory network by integrating known human transcription factor (TF) binding sites with gene expression data in normal human fibroblasts using the algorithm PANDA (33) (see Materials and Methods for further details). After thresholding the edge weights and applying CONDOR (9), a method for community detection in bipartite networks, we found that the baseline network had five communities of varying sizes. Next, we simulated a set of perturbed networks by choosing a random subset of TFs and genes and adding new edges between them, thus artificially creating a new module. The new module consisted of between 3 to 21 TFs, and five times as many genes as transcription factors.

To these simulated networks, we applied three differential community detection methods – community comparison, edge subtraction, and ALPACA – and ranked the nodes by their contribution to the final score for each method. We then used Kolmogorov-Smirnov and Wilcoxon tests to evaluate whether the “true” module ranked higher than expected by chance in each ranked list. The edge subtraction method demonstrated superior performance for recovering modules of all sizes (Figure 2A); this is to be expected, since the only new edges added to the networks were within the new modules. Examining the results from the other two methods, we observed that ALPACA is substantially better than community comparison at detecting smaller modules. Specifically, in a network with a total of ~2500 nodes, community comparison was unable to detect new modules with less than ~110 nodes, whereas ALPACA could reliably detect modules as small as 66 nodes.

**Figure 2.**
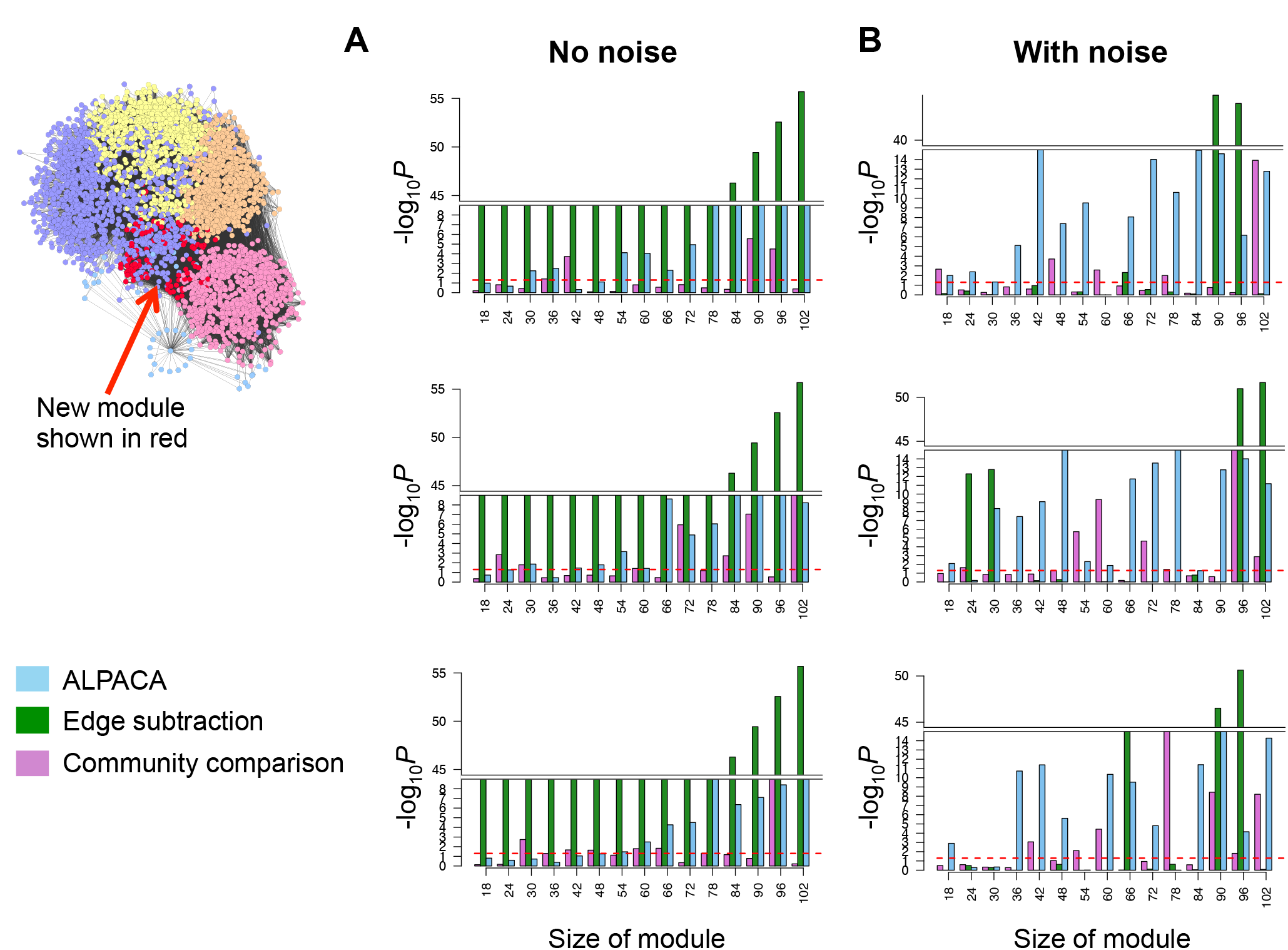
Performance of three methods on simulated networks with added module. Network at left visualizes the regulatory network derived from normal human fibroblasts, with purple, yellow, orange, pink and blue denoting the pre-existing community structure, and red nodes depicting the synthetically added module. Bar graphs show performance of each method – ALPACA, edge subtraction or community comparison – on three random and independent network simulations with **(A)** or without **(B)** resampling of edges among the preexisting communities. P-values computed using Wilcoxon test.

We then introduced edge noise into the “addition” simulation while retaining the modular structure of the underlying network. To do this, we made another series of perturbed networks where, in addition to introducing the new module as described above, we also randomly resampled the edges from the baseline network while retaining the inter- and intra-community edge density. In this more realistic set of simulations, we found that ALPACA outperformed the other methods across a range of module sizes (Figure 2B).

To check that these results are independent of the particular optimization algorithm used, we repeated the analysis using the Louvain method instead of CONDOR for initial community detection in the community comparison and edge subtraction methods. The results were very similar in both cases (Supplementary Figure 1). This indicates that the superior performance of ALPACA is not due to the optimization method used, but rather arises directly from the definition of the differential modularity.

While the edge subtraction method works well to detect “added” modules under low noise conditions, it becomes problematic if edges are deleted or if their weights decrease in the perturbed state relative to the control, because most network clustering methods are only formulated for positive edge weights. One might suggest transformation of edge weights, but any simple transformation of negative edge weights to make them positive (for example, by exponentiation or a linear shift) would bias the results. Algorithms that directly incorporate negative edge weights are complex and involve multiple steps and assumptions (31, 32). In contrast, ALPACA’s differential modularity matrix *D_ij_* contains both negative and positive values, corresponding to areas of decreasing and increasing edge density relative to the baseline network and its community structure. By optimizing over the sum of *D_ij_*, ALPACA incorporates positive and negative changes in edge density in a symmetric fashion.

As a simple demonstration of ALPACA’s ability to detect community structure changes with negative weights, we created “subtracted” simulations in which selected edges in a baseline network are reduced in weight to produce a substantially different perturbed network structure (Figure 3 and Supplementary Figure 2; see Materials and Methods for more details). In Figure 3, for example, the network consists of two dense node groups, A and B, which are more strongly connected together in the baseline condition (edge weight 0.8) than in the perturbed condition (edge weight 0.2). Therefore, the perturbation causes groups A and B to separate and perform distinct functions; intuitively, this means groups A and B characterize the change in modular structure between the two networks. Because the only change in edge weights is the decrease in edges between A and B, the edge subtraction method results in a network with negative edge weights.

**Figure 3.**
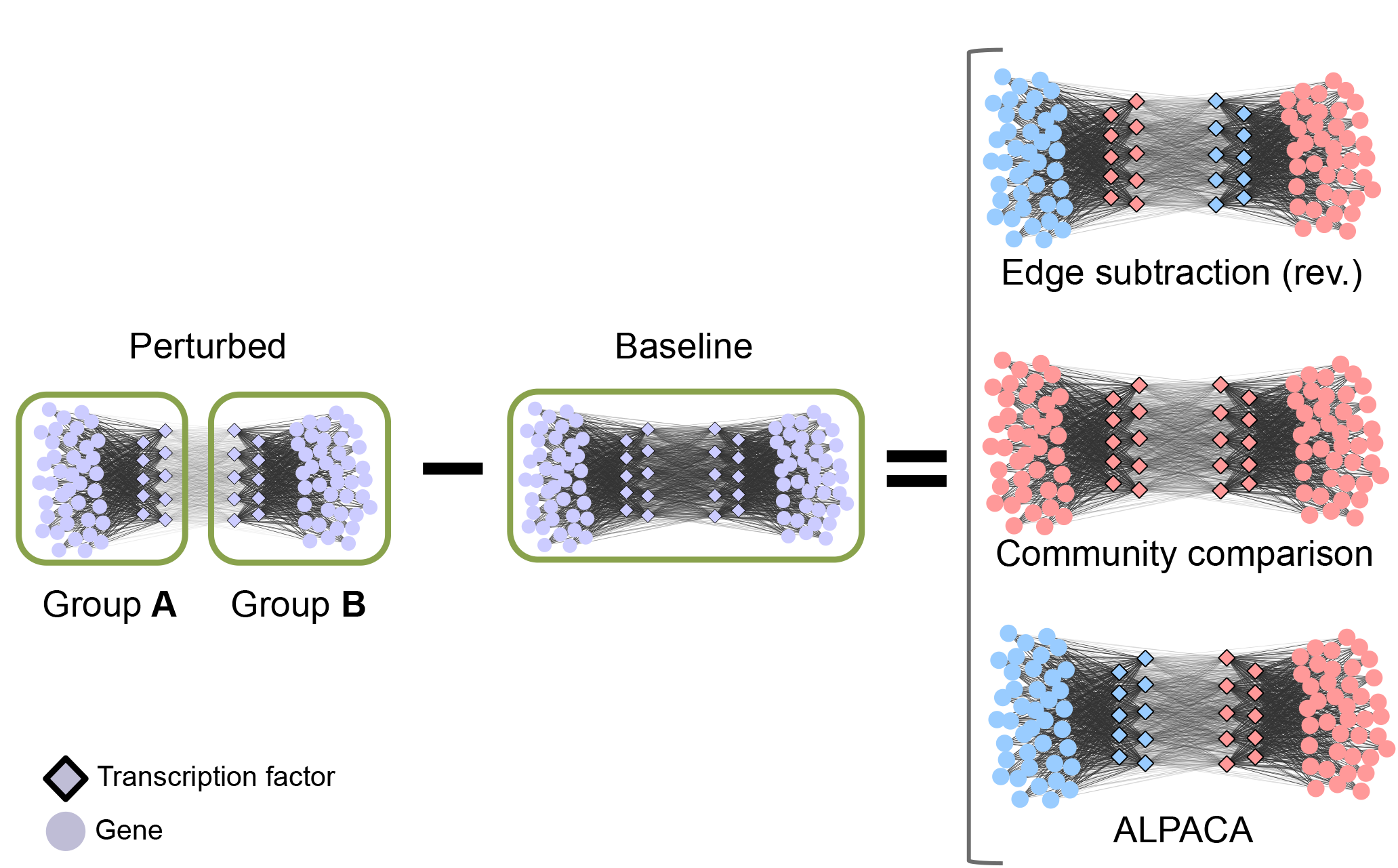
Performance of three methods on perturbations that decrease edge density. Left hand side shows a network transition involving a decrease in edge weights between nodes in Groups A and B. All other edges remain the same. Right hand side shows the results of three methods when comparing these two networks, with the computed differential community structure indicated by node coloring. Note that the “edge subtraction” method needs to be applied in the reverse manner, comparing the baseline network against the perturbed network, in order to have positive differential edge weights.

If instead we reverse the process and subtract the perturbed network from the baseline network, the resulting positive edge weight network produces two modules, one consisting of TFs in group A linked with genes in group B, the other consisting of TFs in group B linked with genes in group A. This does not match the intuitive result we are looking for. The community comparison method detects no change because both the baseline and perturbed networks are composed of the same two node communities. However, ALPACA correctly identifies groups A and B as the differential modules characterizing this transition.

An example with three node groups is shown in Supplementary Figure 2. Again, we find that ALPACA identifies the key change in modular structure and edge subtraction cannot. Although these examples are simple, such areas of decreased edge density will be locally embedded in any realistic biological network and will strongly influence the identification of neighboring modules.

### Angiogenic vs. non-angiogenic ovarian cancer tumors

Ovarian cancer is the second most common cause of cancer death among women in the developed world. Available treatment options for ovarian cancer, such as platinum-based therapies, often lead to chemoresistance and recurrence. Ovarian cancer tumors can be stratified by gene expression profile, tissue of origin, or other characteristics, in order to better understand heterogeneity and predict patient-specific therapeutic strategies. We previously found that a gene signature associated with angiogenesis is able to classify ovarian cancer patients into a poor-prognosis subtype (34).

We classified 510 ovarian cancer patients from The Cancer Genome Atlas into 188 angiogenic and 322 non-angiogenic tumors and used PANDA to infer separate gene regulatory networks for the two subtypes, as described in (35). We then applied a variety of methods to look for changes in community structure associated with the angiogenic tumors, ranked the nodes by their contribution to the total score for each method (see Materials and Methods), and evaluated the core genes in each set for functional enrichment. In order to evaluate the unique contributions of ALPACA, we first applied standard community detection techniques to identify communities in each subtype-specific network, using both the Louvain method and CONDOR, and we looked for GO terms that were statistically enriched in the angiogenic network but not in the non-angiogenic network. Next, we applied edge subtraction, community comparison, and ALPACA to directly identify differential modules associated with angiogenic tumors. The GO term enrichment with *P*_adj_ < 0.05 for each method is presented in full in Supplementary Table 1.

Consistent with what we observed in the simulated networks, ALPACA had higher resolution than the other methods and identified 17 modules specific to the angiogenic network. Strikingly, ALPACA was the only method that identified a gene module enriched in “blood vessel development,” the pathway that we know drives the phenotypic difference between these two ovarian cancer subtypes. Standard community detection methods did not find such a cluster. The non-angiogenic network communities were enriched for histone methylation, embryo development, G-protein coupled receptor signaling, interferon signaling, and chromatin assembly, whereas the angiogenic communities were enriched for cAMP biosynthetic process, response to fibroblast growth factor, MAPKK activity and interferon signaling (Supplementary Table 1). The community comparison method did not yield any enriched GO terms. The edge subtraction method resulted in four large modules enriched for general processes like regulation of cell shape, extracellular matrix organization, nucleosome assembly, and immune response (Supplementary Table 1).

ALPACA led to more specific GO term enrichment than the other methods, suggesting that it was able to more carefully refine differential module structure. For example, instead of general GO terms like “immune response,” the ALPACA modules were enriched for particular immune-related pathways like Type I interferon response, interleukin production, and regulation of the NFκB pathway, and inflammation. Other enriched pathways included JAK-STAT and growth hormone signaling, urogenital development, triglyceride homeostasis, flavonoid glucuronidation, and cell migration. Some of these pathways, like JAK-STAT and cell migration, have already been associated with ovarian tumor progression, while others like flavonoids and triglycerides have only tentative connections with risk of ovarian cancer. We note that most of the ALPACA GO term results could not be found by running community detection on the angiogenic network alone, which shows that ALPACA partitions nodes in a novel manner that does not merely reflect the underlying community structure of the disease network but instead highlights the changes in modular structure between conditions (Figure 4, inset).

**Figure 4.**
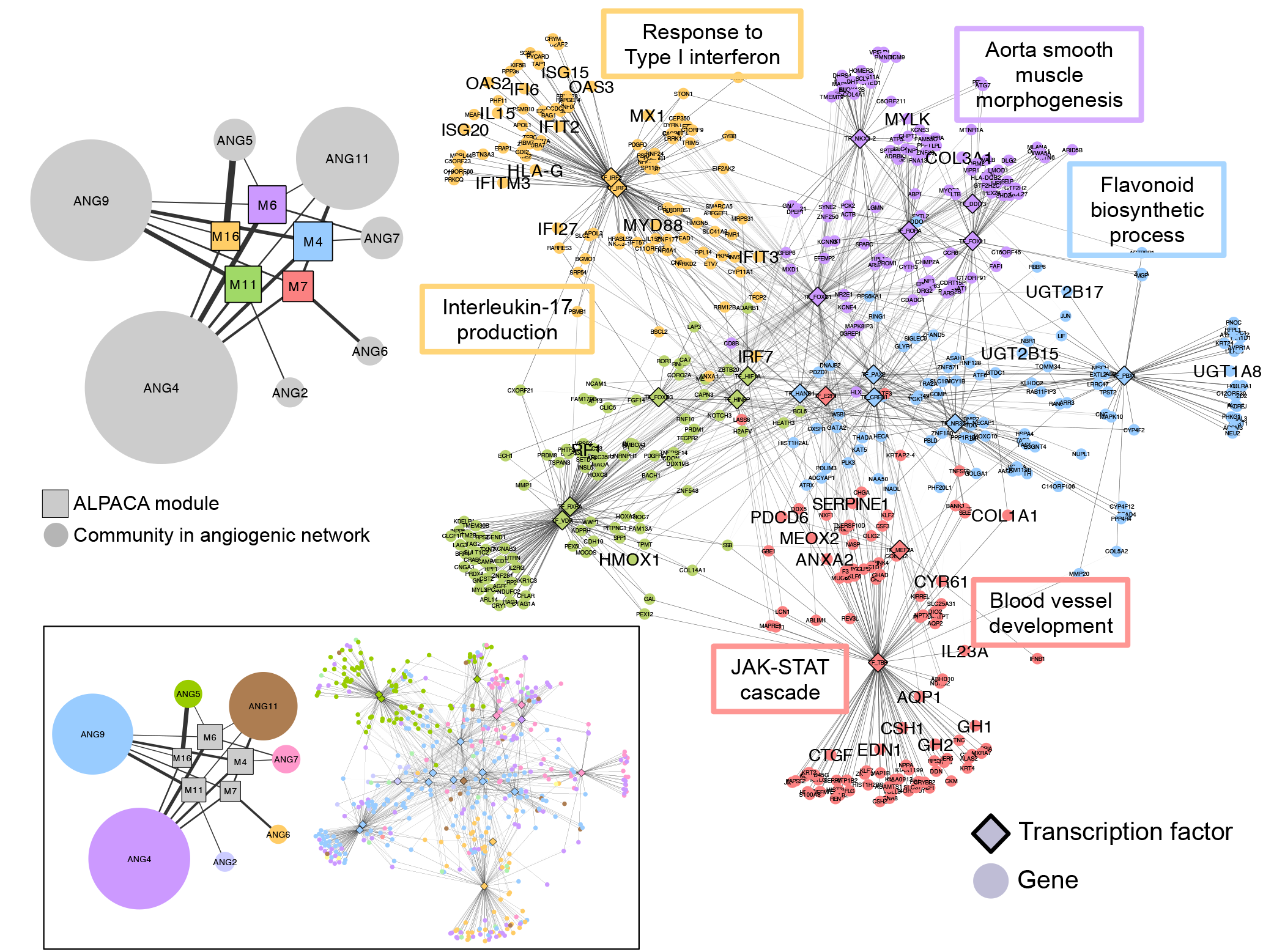
ALPACA modules associated with angiogenic ovarian tumors. Right hand side shows five of the modules, with nodes colored by their membership. Edge opacity is proportional to its contribution to the differential modularity. Network is annotated with representative enriched GO terms with *P*_adj_ < 0.05, and the genes annotated by the shown GO terms are labeled in larger font. Left hand side shows the relationship between the ALPACA modules (denoted by M) and the community structure of the angiogenic network (denoted by ANG). Edge thickness depicts the fraction of genes in that differential module that are present in a particular angiogenic network community. The size of each node is proportional to the number of genes in that module or community. Bottom inset: Same networks as above, but colored by community membership in the angiogenic network rather than by membership in the ALPACA modules.

We also note that running ALPACA in reverse, to find modules present in the non-angiogenic network as compared to the angiogenic network, results in a substantially smaller set of enriched GO terms, which fall mostly into the metabolic and immune categories, with no enrichment in blood vessel development (Supplementary Table 1). ALPACA therefore selectively identifies biological signals associated with the specific phenotype under study.

We examined the ALPACA modules and their connections to ovarian cancer in more detail, focusing on non-redundant GO terms that had an overlap of three genes or more with the module in which it was enriched (Figure 4). Module 4 was enriched for “flavonoid glucuronidation” and contains the UDP glucuronosyltransferases UGT2B15, UGT1A8, and UGT2B17, enzymes that can help metabolize flavonoids and regulate hormones. Studies have hinted that dietary intake of flavonoids may reduce the risk of ovarian cancer (36–38) but the association is not statistically robust, and the mechanism is unknown. Our results suggest that the UGT family of enzymes may mediate the connection between flavonoids and ovarian cancer. Module 5 is enriched in “urogenital system development” and contains several genes that are highly relevant to ovarian cancer. HNF1B is known to be a subtype-specific ovarian cancer susceptibility gene (39). Its expression level and promoter methylation status is predictive of clear cell and invasive serous subtypes of epithelial ovarian cancer. ESR1 is the estrogen receptor and is central to breast and ovarian cancer. IQGAP1 is a scaffold protein whose expression appears to drive invasion and progression of ovarian cancer tumors (40–42). SOX11 acts as a tumor suppressor in ovarian cancer, and its expression is regulated by methylation and predicts patient survival (43, 44).

We found that module 12 was enriched in triglyceride homeostasis. Although it is not known whether there is a dietary effect of triglycerides on ovarian cancer risk, several studies have noted that ovarian carcinomas have distinctive lipid profiles and metabolic characteristics (45). Our results suggest that metabolic pathways involving hepatic lipase C (LIPC) and glucokinase regulator (GCKR) may be mobilized differently in poor-prognosis ovarian cancer. Finally, modules 16 and 17 were enriched for various terms involving interferon response, interleukins, and regulation of the NFκB pathway, consistent with the theory that chronic inflammation is associated with risk of cancer (46). Specifically, the interleukin IL6 has been proposed as a therapeutic target, and IL12 is a prognostic factor in ovarian cancer (47–49). Interferons have cytotoxic properties in ovarian cancer cells (50, 51). NFκB activation is correlated with poor prognosis in ovarian cancer, and blocking the NFκB pathway can reduce anchorage-independent growth and invasiveness in cell culture assays (52).

Module 7 was enriched in “blood vessel development” and “positive regulation of cell migration,” reflecting the invasive and angiogenic characteristics of poor-prognosis ovarian tumors. The apoptosis gene PDCD6 is a member of both GO terms and is topologically central to this module. Interestingly, it is a known predictor of progression free survival in ovarian cancer and synergizes with cisplatin to inhibit ovarian cancer cells *in vitro* (53–55). CYR61, also a member of both GO terms, is an extracellular matrix (ECM) signaling protein that is overexpressed in poor prognosis ovarian carcinoma (56, 57). CTGF (connective tissue growth factor) is an angiogenic ECM protein, and it appears to have an inverse relationship with CYR61; high CTGF expression correlates with low CYR61 in low-grade tumors with increased survival (58). Overall, this suggests that the ALPACA modules contain functional groups of prognostic genes that may interact with each other to produce distinct phenotypes. ALPACA could therefore be a useful feature selection step to isolate small groups of pathway genes and build more complex predictive models.

Module 7 was also enriched in growth hormones and the JAK-STAT cascade. The JAK-STAT pathway is constitutively active in breast, ovarian and prostate cancers, and nuclear localization of activated STAT3 is associated with worse survival and chemoresistance in ovarian cancer. Treatment with JAK2 inhibitor reduces tumor burden in ovarian cancer xenografts (59). Members of module 7 that are annotated with this GO term include growth hormones 1 and 2 (GH1 and GH2). This pathway is already known to be a drug target in ovarian cancer, and growth hormone-releasing hormone (GHRH) antagonists reduce proliferation of ovarian cancer cells both *in vitro* and *in vivo* (60–62).

### Tumor virus perturbations in primary human cells

DNA viruses hijack the host cell cycle to jumpstart viral genome replication. Tumor viruses can do this so effectively that they lead to aberrant cell proliferation and tumorigenesis, and studying tumor viruses can shed light on the molecular mechanisms behind cancer. Previously, we expressed a panel of 63 proteins from four families of DNA tumor viruses – Epstein-Barr virus (EBV), human papillomaviruses (HPV), polyomaviruses, and adenovirus – in IMR90 primary human fibroblasts and generated gene expression profiles for each cell line (63). To construct regulatory networks, we divided the gene expression data into two groups, the first corresponding to the 37 viral proteins classified as “transforming” due to their tumorigenic properties, and the second corresponding to all the control cell lines that contain either empty vectors or GFP. We used PANDA to infer networks by combining gene expression from each sample group with a prior map of cell type-specific DNase-I-hypersensitive TF binding sites (33).

We first ran standard community detection on each network, using the Louvain method for modularity maximization. The control network contained communities enriched in cell migration, axon guidance, and wound response (Supplementary Table 2). The communities in the transforming viral oncogene network were enriched for epithelial-mesenchymal transition, cell migration, axon guidance and wound response. Since an important function of fibroblasts is to migrate and heal wounds, many of the results from standard community detection appear to be cell type-specific processes that are not specific to viral oncogenes. The genes with the biggest changes in community assignment were enriched in BMP response and natural killer cell development (Supplementary Table 2). Applying the edge subtraction method using Louvain or CONDOR optimization methods resulted in enrichment for chromatin modification, the Toll-like Receptor (TLR) pathway, and immune response. We then applied ALPACA to compare the two networks. Like the edge subtraction method, ALPACA also revealed changes in immune response and chromatin modification but, importantly, it also found significant enrichment for “mitotic cell cycle,” which is the main process we expect to be perturbed by tumor viruses (Figure 5). Consistent with this, we had previously found that fibroblast cell lines expressing transforming viral oncogenes have significantly altered growth rates (63).

**Figure 5.**
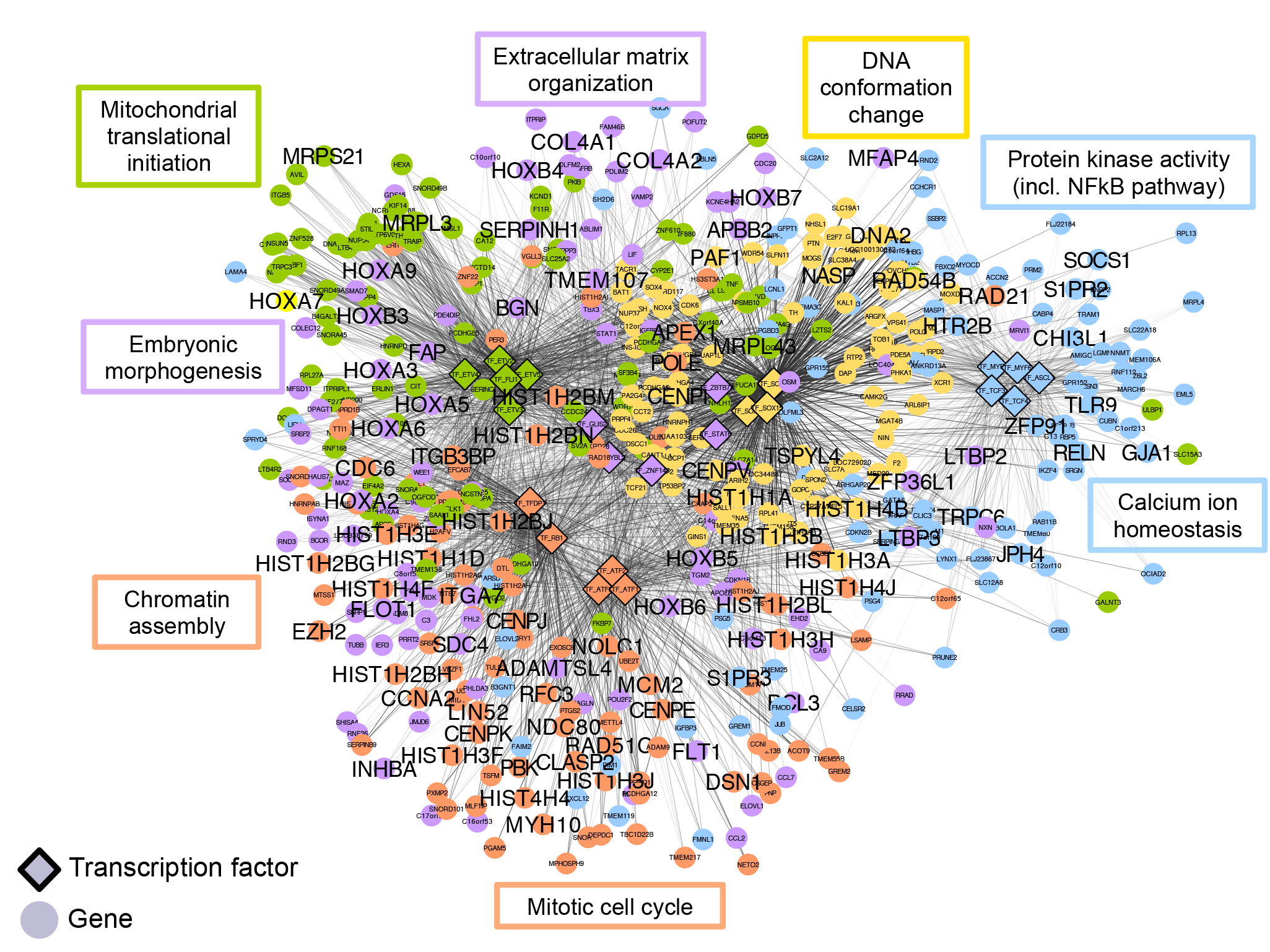
ALPACA modules associated with transforming viral oncogenes. Network shows five modules, with nodes colored by membership in differential modules. Edge opacity is proportional to its contribution to the differential modularity. Network is annotated with representative enriched GO terms with *P*_adj_ < 0.05. Genes annotated by the shown GO terms are labeled in large font.

ALPACA was also the only method to identify communities representing several cancer pathways that are known to be targeted by tumor viruses, including extracellular matrix (ECM) organization, NFκB signaling, and embryonic development. Module 1 is enriched in “cellular calcium ion homeostasis” and “regulation of NIK/NF-kappaB signaling.” NFκB and Nuclear factor of activated T-cells (NFAT) are two important cancer-related pathways that activate immune cells, and NFAT activity is modulated primarily through intracellular calcium levels. Merkel cell polyomavirus and EBV LMP1 are both known to functionally perturb the NFκB pathway through different mechanisms (64, 65). EBV, HPV16 and several polyomaviruses target the genes CHI3L1, TLR9, and SOCS1, which are all among the top-scoring nodes in this module (66–71). EBV and HPV infections both alter calcium signaling in the host cell (72, 73). Tumor viruses use these pathways in a variety of ways to increase cell growth and manipulate the innate immune response.

We found that module 4 was enriched in many terms related to “embryonic morphogenesis” and development. We previously found that tumor viruses co-opt the Notch pathway, which is central to embryonic development, in order to promote cell growth and tumorigenesis (63). The GO term enrichment among the target genes in module 4 is driven by the homeobox (HOX) TFs, whose expression is regulated by EBV LMP1 and HPV E7 through differential methylation (74–76). Module 4 was also enriched in “extracellular matrix organization.” The epithelial-to-mesenchymal (EMT) transition is a key step in epithelial tumorigenesis, and cells undergoing EMT often acquire the ability to degrade extracellular matrix (ECM) proteins and increase their invasive potential (77). In particular, the transforming HPV E6 and E7 proteins are able to upregulate matrix metalloproteinases (MMPs) in order to degrade ECM and increase cell migration, thus leading to cellular transformation (78).

Both ALPACA and the edge subtraction method detected a difference in the regulation of histones, suggesting that epigenetic changes may be a key factor in the transformation of human cells by viral oncogenes. ALPACA also identified a separate module (module 8) that was enriched in proteins involved in “DNA conformation change.” Indeed, the importance of epigenetics in transformation has already been demonstrated for many tumor viruses. HPV16 E7 induces histone 3 lysine 27-specific demethylases (76), EBV LMP1 and LMP2A modulate the activity of DNA methyltransferases and interact with histone modifiers (79), and adenovirus E1A causes sweeping changes in histone acetylation (80).

### Sexual dimorphism in normal breast tissue

The Genotype-Tissue Expression (GTEx) consortium has generated gene expression data using tissue collected from 51 body sites and in nearly 600 individuals. Not surprisingly, the tissue with the greatest difference between males and females in autosomal gene expression is the breast (81). We used PANDA to create tissue-specific regulatory networks to study the effect of sex on regulatory networks in breast tissue (81). We first applied the Louvain method to detect communities separately in the networks derived from male and female breast tissue and tested for functional enrichment of GO terms in the male and female communities. We found that both the networks were enriched for the same biological processes: GTPase-mediated signal transduction and protein catabolic process (Supplementary Table 3). Therefore, despite what one might expect to be substantially different, the global structure of the male and female networks failed to identify sex-specific patterns of regulation. We also used the edge subtraction method to search for modular differences between the sexes and tested modules for GO term enrichment, but this too failed to identify any significant GO biological processes.

We then tested whether ALPACA could find sex-specific modular structure in the breast regulatory network (Figure 6). We first compared the male regulatory network against the female regulatory network and found 18 male-specific differential modules (Supplementary Table 3). Module 2 was highly enriched in developmental processes, including “nervous system development,” “response to BMP,” and “blood vessel development.” Similarly, module 8 was enriched for “muscle organ morphogenesis.” These results are not surprising and reflect the fact that male and female breast tissues have significant differences in their developmental trajectory. We note that many of the developmental genes in these modules are associated with breast cancer. Among genes annotated with “nervous system development,” the fibroblast growth factor receptor (FGFR) is often amplified or dysregulated in breast cancer, the HES5 locus is repositioned in invasive breast cancer, and VLDLR is often upregulated in metastatic breast cancer (82–85). The blood vessel development category included genes such as GATA6, a known oncogene that may drive EMT in the breast; TBX3, which appears to repress the tumor suppressor p14ARF and drive metastatic breast cancer; PRRX2, which increases invasiveness in breast tumors; and RASA1, whose expression is associated with poor prognosis in breast cancer (86–89). Among the BMP response and muscle development groups, there are several genes, like TWSG1, VANGL2, and GSC, which are relevant to both normal breast development and breast cancer (90–92). Module 7 was enriched for terms related to rRNA processing and module 14 contained genes relevant to chromatin assembly, suggesting that transcription and translation are reorganized at a global level between males and females.

**Figure 6.**
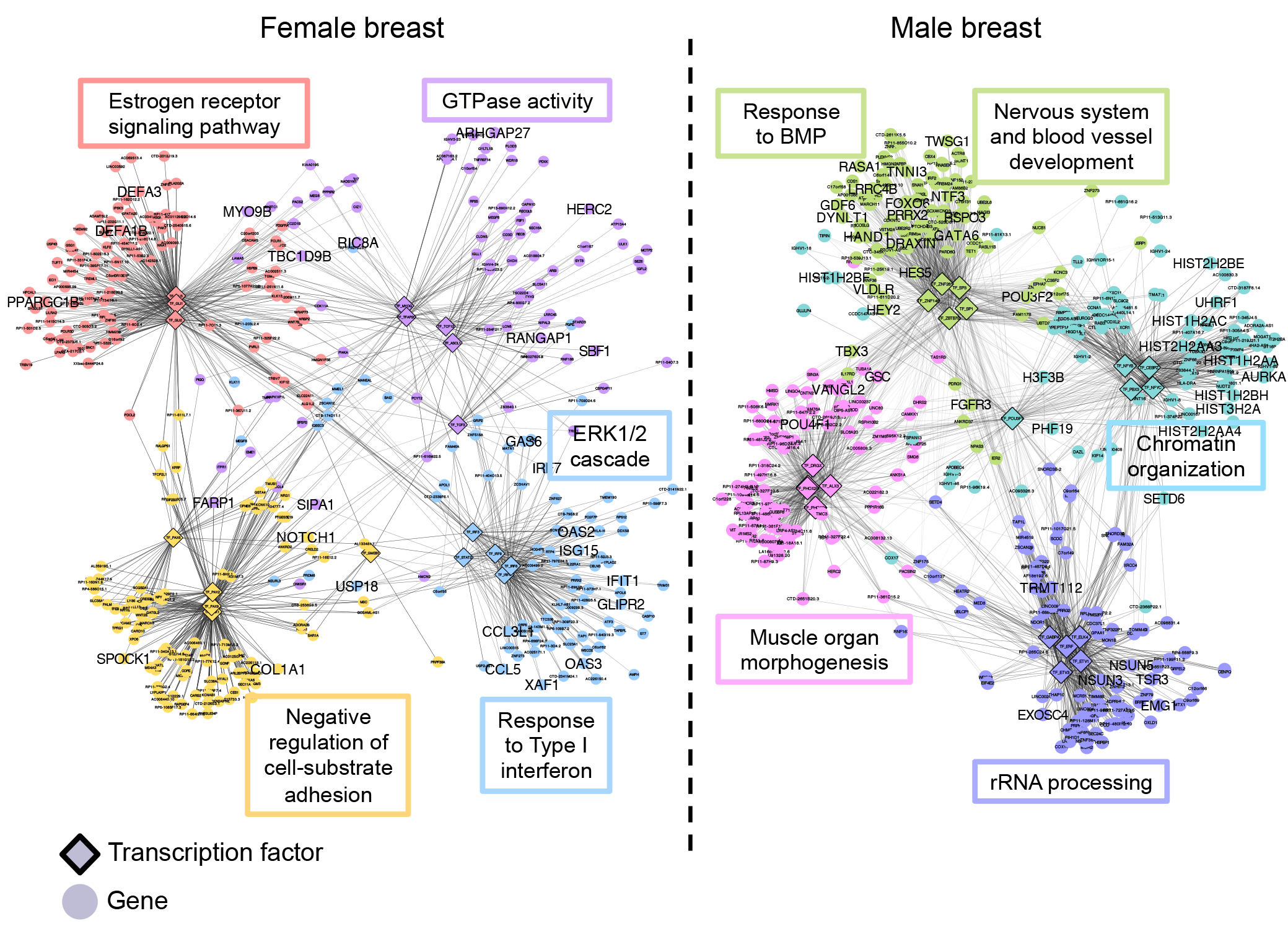
Sexually dimorphic ALPACA modules in human breast tissue. Networks show four modules specific to either female (left-hand side) or male (right-hand side) breast tissue. Nodes are colored by membership in differential modules. Edge opacity is proportional to its contribution to the differential modularity. Networks are annotated with representative enriched GO terms with *P*_adj_ < 0.05. Genes annotated by the shown GO terms are labeled in large font.

Next, we compared the female breast regulatory network against the male network and found 17 female-specific regulatory modules. Among those, ALPACA identified a module (module 15) that is enriched in “intracellular estrogen receptor signaling pathway,” the hormonal process one would expect to be critical for female breast development and overall function. The highest-scoring gene in this pathway, PPARGC1B, is a co-activator of the estrogen receptor and is a genetic risk factor for estrogen receptor-positive breast cancer (93, 94). Module 10 was enriched for “positive regulation of ERK1 and ERK2 cascade.” ERK1/2 signaling is a major pathway involved in estrogen-induced cell proliferation and breast cancer (95, 96). This module contains the growth arrest-specific gene GAS6, which is induced by estrogen and is associated with chemoresistance and metastasis in breast cancer (97–99), and the chemokine CCL5, which has been proposed as a therapeutic target for estrogen-dependent breast cancer (100, 101). Module 10 was also enriched for Type I interferon response, which may be a result of the increased blood and lymphatic penetrance in normal female breast development.

We found module 17 to be enriched for “negative regulation of cell-substrate adhesion” and contained SPOCK1 and NOTCH1, both known markers of invasion and breast cancer progression (102–105). Finally, module 5 was enriched in transcriptional regulation factors, similar to the enrichment in chromatin remodeling found in the male breast network.

Consistent with expectations based on both the functional differences of male and female breast, and the profound differences in gene expression, ALPACA was able to identify major biological processes associated with differences in breast development between females and males, many of which are also known to be dysregulated in breast cancer.

## DISCUSSION

Biological networks have complex modular and hierarchical topologies that allow organisms to carry out the functions necessary for survival. Various perturbations, such as environmental conditions or mutations, can alter regulatory networks, leading to changes in the phenotype of the organism. Techniques such as differential expression analysis can be used to characterize the transition between different cellular states, but changes in gene expression are ultimately driven by changes in regulatory pathways. If we are to build predictive models of complex phenotypes and diseases, it is essential that we understand how the regulatory network also changes with phenotype. ALPACA is an algorithm that compares genome-scale networks using a metric we call the “differential modularity” to find groups of nodes that drive changes in modular structure. ALPACA differs from other network methods in that it compares the structure of networks to each other rather than to a random background network and is thus more likely to detect subtle differences in network modular structure. This potentially can allow detection of gene modules that function together in particular conditions or in disease.

We evaluated the performance of ALPACA on simulated networks and compared it to two available approaches for detecting changes in network modular structure: (i) “community comparison,” where one applies standard community detection to the baseline and perturbed networks separately and contrasts the resulting communities, and (ii) “edge subtraction,” which involves subtracting the two networks edge by edge, and clustering the resulting differential network. ALPACA was able resolve smaller differential modules than the community comparison method. Intuitively, this is because modularity maximization in its standard form penalizes the splitting of a large dense community into smaller ones, whereas the differential modularity score used in ALPACA penalizes the formation of large communities similar to those present in the baseline network. In addition, ALPACA was more robust to noise in individual network edges than the edge subtraction method. In the edge subtraction method, the uncertainty of the edges in the “differential” network is the sum of the uncertainties in the corresponding edges of the original networks. Instead, ALPACA aggregates the signals coming from multiple edges in the baseline network communities to derive a null model for edge density, so it is less sensitive to the uncertainty in individual edges.

ALPACA’s differential modularity metric directly compares the edges that one sees within a community to what you would expect based on the topology of a corresponding reference network. This adapts the well-established modularity maximization method to infer subtle changes in the community structure that arise when comparing distinct complex phenotypes. Unlike other methods that simply subtract networks, ALPACA preserves those secondary interactions that exist in both networks but allows them to shift their functional context as the edges around them change, which can capture new modular structures. The differential modularity also incorporates increased and decreased edge weights across the entire network into a single, simple framework for module detection. And unlike community comparison, ALPACA can detect new modules that form on top of a background of globally active regulatory programs that are present in both the baseline and perturbed networks.

We applied ALPACA to transcriptional networks that were constructed from gene expression and TF binding data using the PANDA network inference algorithm. PANDA does not explicitly use the expression correlation between regulators and the target genes, and can therefore model TFs that are not changing in mRNA expression but whose activity is controlled through other mechanisms, like post-translational modification. PANDA also incorporates changes in promoter activity that could alter regulatory targeting patterns. Comparing angiogenic to non-angiogenic subtypes of ovarian cancer, we found functional modules that were enriched in expected disease pathways like blood vessel development, interleukin production, and JAK-STAT signaling. We also found enrichment for less expected processes including nutritional pathways like flavonoid biosynthesis and triglyceride homeostasis, which have been speculated to be relevant for ovarian cancer, but for which the underlying molecular pathways are not known (36–38, 45). These biological processes were specific to the angiogenic subtype and uniquely revealed by ALPACA; they could not be found through standard community detection in the individual angiogenic and non-angiogenic networks or in an edge-subtracted network, or by running ALPACA in reverse on the non-angiogenic network compared to the angiogenic network.

In another test of the method, we compared normal male and female breast tissue to find sex-specific patterns of regulation. Many of the modules we found were enriched in known processes related to breast development and breast cancer, like ERK and Rho GTPase signaling. Perhaps most strikingly, the female breast network contained a differential module enriched for estrogen receptor signaling, which is one of the main sex-specific pathways known to be active in breast tissue. Once again, these results could not be found using other community detection and network comparison methods.

ALPACA requires a minimum input of two graphs. It is easily generalizable and could be applied to many types of biological networks, including metabolic, protein-protein interaction (PPI), and expression Quantitative Trait Loci (eQTL) networks, all of which exhibit highly functional modular structures (9, 106, 107). For example, we could imagine applying ALPACA to compare community structure in PPI networks with mutation-driven “edgetic” perturbations, in order to discover functional changes in protein complexes and signaling associated with disease (108). ALPACA could also be applied to compare eQTL networks in patient cohorts with differing pathologies to prioritize sets of SNPs and genes that influence complex traits (9).

ALPACA builds on our growing understanding of how networks define phenotype. Differential expression is driven by changes in the activity and structure of gene regulatory networks. But adding or subtracting edges does more than change individual regulatory interactions. With enough individual changes occurring in the right places in the starting network, changes in edges can lead to the creation or destruction of functional communities of genes and their regulators. While the global structure of the network may be largely unchanged, these new functional communities provide insight into coherent processes that differentiate one phenotype from another.

As more genome-wide studies of molecular interactions and multi-omics data are generated, better statistical models for network analysis will be critical to making differential network biology a robust and reproducible platform for studying complex diseases (13). To transform this large volume of data into clinically useful predictions and hypotheses, we need rigorous methods that can integrate heterogeneous data types and extract the functional elements that are key parameters for modeling disease transitions and the genotype-phenotype relationship. ALPACA is the first method to make direct comparisons between networks to identify changes in their modular structure in a rigorous manner and is an important step forward in methodology in the statistical analysis of networks.

## MATERIALS AND METHODS

### ALPACA algorithm

ALPACA is implemented in R and is freely available for download through Github at https://github.com/meghapadi/ALPACA. It is comprised of the following two steps:

*Step 1:* The input network consists of edges between regulators and target genes. We first label the nodes that act as regulators and targets separately. In particular, a gene that encodes a transcription factor (TF) becomes two separate nodes depending on whether we are modeling its mRNA expression level (target node) or protein activity (regulator node). For the weight of each edge, we use the final z-score output by the PANDA network inference algorithm. We then take the edges that have positive weight in the baseline condition, and run bipartite weighted network community detection using either CONDOR or the Louvain method.
*Step 2:* Compute *D_ij_* for the perturbed network, using the definition in the main text and the baseline communities found in Step 1. It is possible that the numerator and denominator of *N_ij_* are both zero, meaning that there were no edges between the communities *C_i_* and *C_j_*. This can happen if, for example, at least one of the nodes *i* or *j* were not connected to the baseline network to begin with. In this case, we define *N_ij_* to be zero, since the “expected” number of edges between the two nodes is zero. We next apply a generalized Louvain procedure to assign nodes into communities based on *D_ij_* (18). Briefly, the Louvain method works as follows: (i) Start with every node in its own community, (ii) go through each node iteratively, and merge it with the node that produces the biggest increase in differential modularity, (iii) after reaching a local optimum, treat each of the resulting groups as “metanodes” in a new “metanetwork” and recalculate an effective adjacency matrix, and (iv) repeat steps (ii) and (iii) until convergence. For the purpose of reporting reproducible results, we iterate through the nodes in the same pre-determined order every time, and we break ties by selecting the first member of the set. In an optional third step, we can evaluate the core genes in each module for enrichment in known biological pathways.
*Step 3:* The core genes are those that are most important to the integrity of the module and therefore potentially the most robust and essential members. To define the core genes, we score each node according to its contribution to the differential modularity of the module that it belongs to:

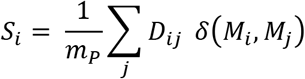

We ranked the target genes in each module by their scores *S_i_*. Since the size of typical modules found in ALPACA ranged from about 50 to 200 genes, we chose to use the top 50 core genes from each module to evaluate functional enrichment in an equitable manner across all the modules. We also repeated each analysis using the top 100 core genes in order to test the dependence of the enrichment on the cutoff. GO term enrichment was calculated using the GOstats package in R, with the following parameters: the gene universe is defined to be the set of all possible target genes in the initial networks, and the p-value calculation is conditioned on the GO hierarchy structure. In each module, the p-values were adjusted for multiple testing using the Benjamini-Hochberg method.

### Edge subtraction method

For each edge, the edge weight of the baseline network was subtracted from the edge weight in the perturbed network to compute *Δw_ij_*, and only edges with *Δw_ij_* > 0 were retained. We then used the *Δw_ij_* values as new edge weights to perform community detection using CONDOR or Louvain optimization (9, 30).

### Community comparison method

We first used either CONDOR or Louvain method to find the community structure of the baseline and perturbed networks, in each case keeping only edges that had positive z-scores. We next aimed to efficiently map the two community structures to each other. To find the best approximation of a linear mapping, we computed *R* in the equation *B* = *AR*, where *A* is the *N x q* matrix of node membership for the baseline community structure, and *B* is the corresponding matrix for the perturbed community structure (here *N* is the number of genes and *q* is the number of communities). To invert the matrix *A*, we used singular value decomposition to compute the pseudoinverse *A^P^* = *VD*^−1^*U^T^*, where *A* = *UDV^T^*, and then computed *R* = *A^P^B*. The entries of the *q* × *q* matrix *R* represent an approximate linear transformation that maps the communities in the baseline network to the communities of the perturbed network. Finally, we scored each node according to how much its community membership remains the same between baseline and perturbed conditions, using the formula 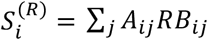. Nodes were ranked from low to high values of 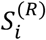 for further analysis. Low-scoring nodes represent the nodes that participate in altered community structure in the perturbed network.

### Creating simulated networks and evaluating differential community methods

To simulate “addition” networks, we started with the GFP-control network from the tumor virus dataset (see section on “Data preprocessing and network inference” for details on how this network was constructed) and thresholded the edges at a z-score of 2.7 (for noiseless simulation) or 2.9 (for noisy simulation). The threshold was chosen such that the resulting edges would form an unweighted network with a similar community structure as the full weighted network. We found that applying CONDOR to the GFP-control network at a threshold of 2.7 resulted in five communities containing 1336, 833, 781, 1018, and 44 nodes each. To add a module, we randomly chose a subset of these nodes and added all possible new edges between them. To add noise in the second set of “addition” networks, we also resampled edges as follows: (i) start with an empty network with the same nodes as the GFP-control network, (ii) count the number of edges between TFs in community *C_i_* and target genes in community *C_j_*, for each pair *i* and *j*, in the GFP-control network, and (iii) add a matching number of edges randomly between the TFs in community *C_i_* and target genes in community *C_j_* in the new network.

We evaluated the results of each method on the simulated networks by comparing the ranks of true positives (the target genes in the added module) against a background consisting of target genes not in the added module. We used Kolmogorov-Smirnov and Wilcoxon tests to look for significant differences in the distribution of the ranks. Both tests gave similar results, and in the figures we present the Wilcoxon p-values.

To create the “subtracted” simulation with two node groups, we started with a fully connected network containing 100 nodes, with all edge weights set to a default value of 0.1. We then defined two node groups, A and B, each containing 10 TFs and 40 genes. Edges within each of these groups were set to edge weight 1.0.

Next, to create the baseline network we set the weights of all edges between groups A and B to be 0.8. To create the perturbed network we set the weights of all edges between groups A and B to be 0.2. To create the three-group “subtracted” network, we first created a fully connected network containing 125 nodes, with all edge weights set to a default value of 0.1. We then defined three node groups A, B, and C containing 50, 25, and 50 nodes respectively (of which 10, 5, and 10 were TFs). Edges within each group were set to weight 1.0, and all edges between groups B and C were set to weight 0.2. For the baseline network, the edges between groups A and B were set to weight 0.8 and for the perturbed network, the edges between groups A and B were set to weight 0.2.

### Data preprocessing and network inference

Preprocessing and network inference for ovarian cancer data was carried out as described in (35). Briefly, we ran the network inference algorithm PANDA (Passing Attributes between Networks for Data Assimilation) to integrate gene expression data with transcription factor binding sites to create regulatory networks for each subtype (33). The prior network of binding sites for 111 TFs were defined as the occurrence of the corresponding motif in the promoter, defined as [−750,+250] base pairs around the transcription start site (TSS).

The viral oncogene gene expression data were normalized and batch-corrected, and a map of high-probability TF binding sites was created by combining cell-type-specific DNase-I hypersensitivity data with motif occurrence in the promoters defined as [-25kb, 25kb] around each TSS, as described in (63). The binding sites and gene expression were combined to infer networks using PANDA with default parameters, as described in (1).

Sex-specific and tissue-specific transcriptional networks for the GTEx data were constructed as described in (81).

## AUTHOR CONTRIBUTIONS

MP conceived of the project, performed analysis, and wrote the paper. JQ helped refine the analyses and write the paper.

## ACKNOWLEDGEMENTS

This work was supported by NIH grants K25 HG006031 (MP) and R01 HL111759 and R35 CA197449 (JQ).

---

**Supplementary Figure 1.**
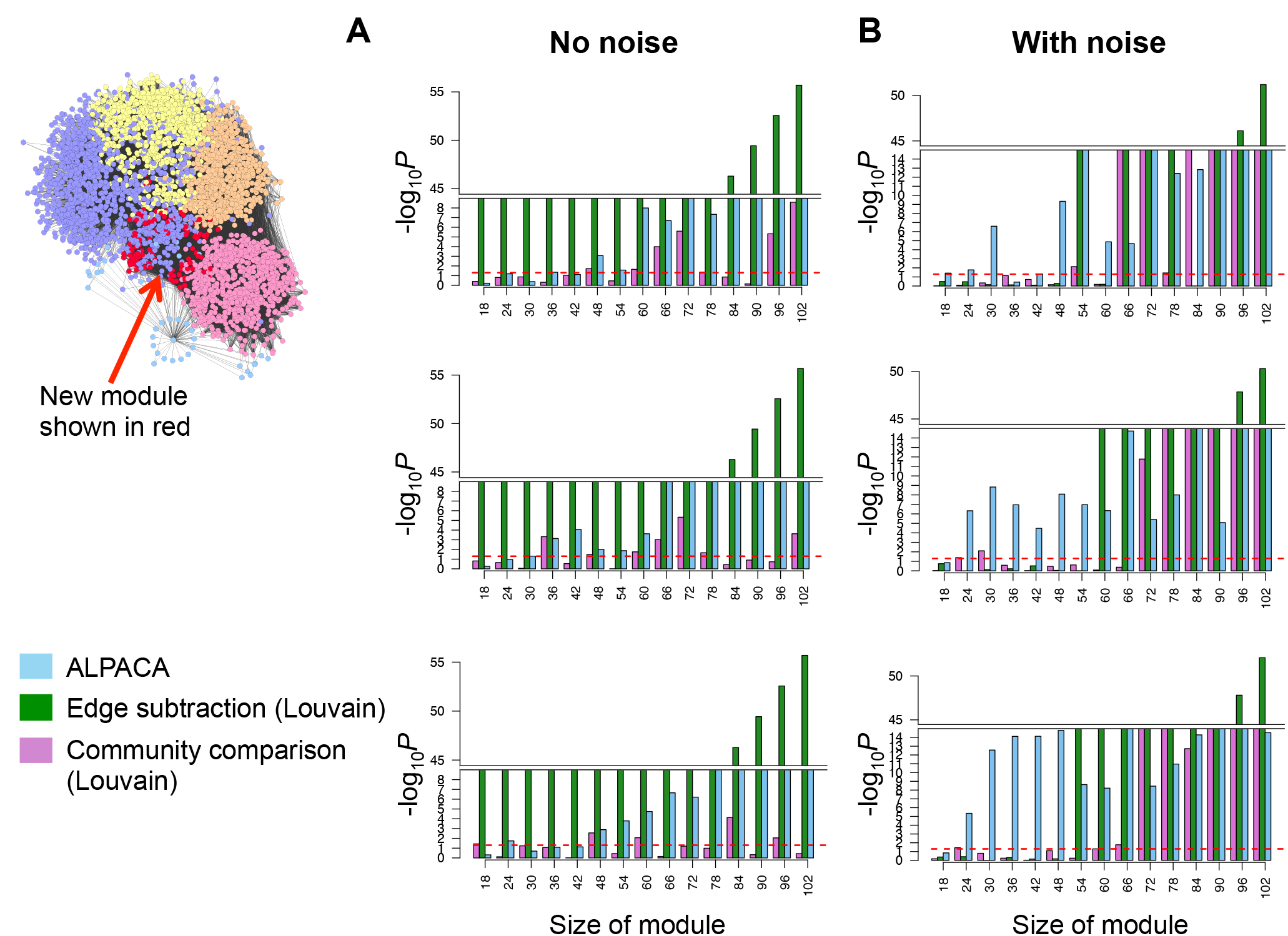
Performance of three Louvain-based differential community methods on simulated networks with added module. Network at left visualizes the regulatory network derived from normal human fibroblasts (as shown in Figure 2). Bar graphs show performance of each method – ALPACA, edge subtraction with Louvain optimization, or community comparison with Louvain optimization – on three random and independent network simulations with **(A)** or without **(B)** resampling of edges among the preexisting communities. P-values computed using Wilcoxon test.

**Supplementary Figure 2.**
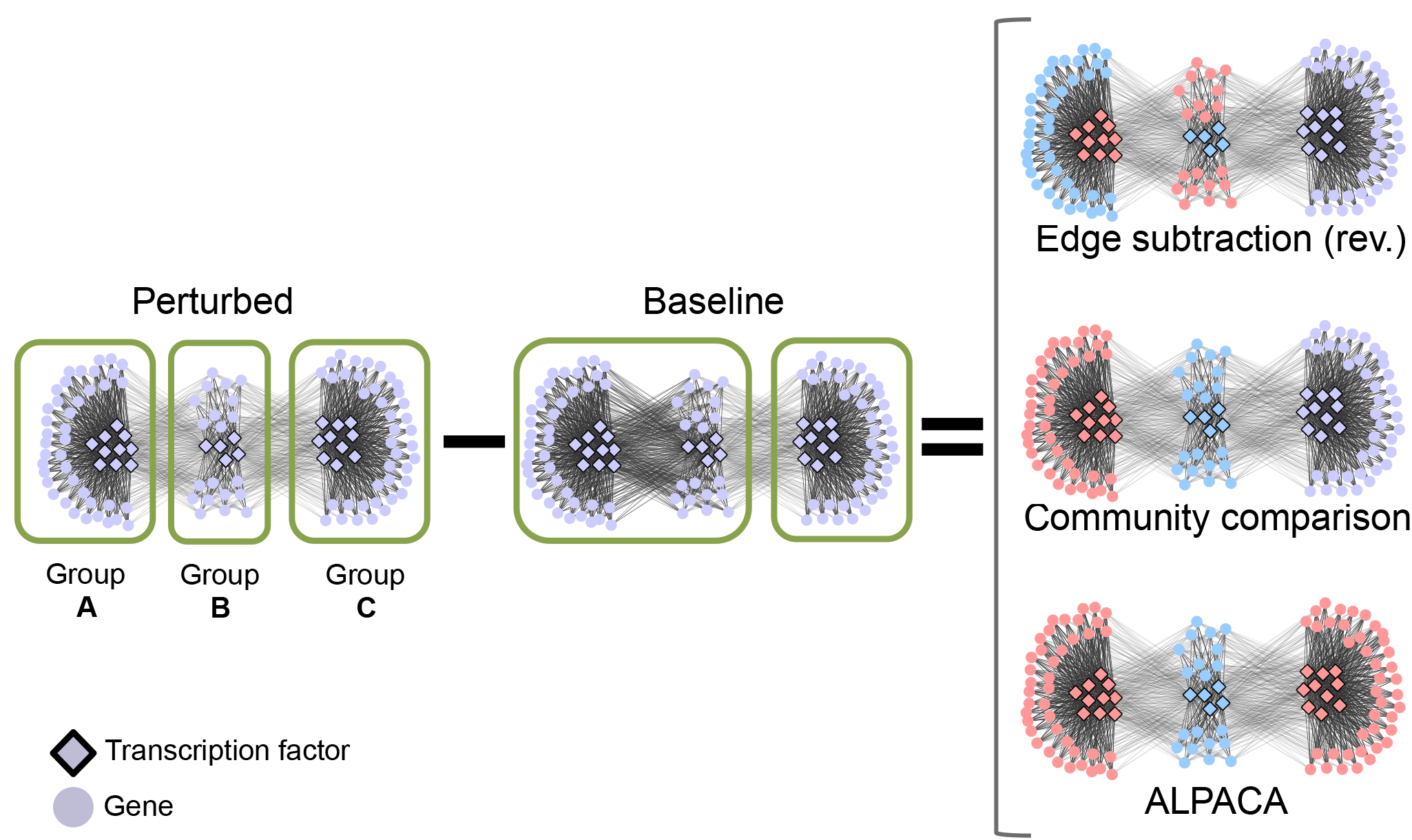
Performance of three methods on three-component network with decrease in edge density. Left hand side shows a network transition involving a decrease in edge weights between nodes in Groups A and B. All other edges remain the same. Right hand side shows the results of all three methods when comparing these two networks. Note that the “edge subtraction” method needs to be applied in the reverse manner, comparing the baseline network against the perturbed network, in order to have positive differential edge weights. The light violet color denotes nodes that remain unclassified by the indicated differential community methods.

